# Cancer-driving mutations are enriched in genic regions intolerant to germline variation

**DOI:** 10.1101/2022.01.07.475416

**Authors:** Dimitrios Vitsios, Ryan S. Dhindsa, Jonathan Mitchell, Dorota Matelska, Zoe Zou, Joshua Armenia, Quanli Wang, Ben Sidders, Andrew R. Harper, Slavé Petrovski

## Abstract

Large reference datasets of protein-coding variation in human populations have allowed us to determine which genes and genic sub-regions are intolerant to germline genetic variation. There is also a growing number of genes implicated in severe Mendelian diseases that overlap with genes implicated in cancer. Here, we hypothesized that mitotically mutable genic sub-regions that are intolerant to germline variation are enriched for cancer-driving mutations. We introduce a new metric, OncMTR, which uses 125,748 exomes in the gnomAD database to identify genic sub-regions intolerant to germline variation but enriched for hematologic somatic variants. We demonstrate that OncMTR can significantly predict driver mutations implicated in hematologic malignancies. Divergent OncMTR regions were enriched for cancer-relevant protein domains, and overlaying OncMTR scores on protein structures identified functionally important protein residues. Finally, we performed a rare variant, gene-based collapsing analysis on an independent set of 394,694 exomes from the UK Biobank and find that OncMTR dramatically improves genetic signals for hematologic malignancies. Our web app enables easy visualization of OncMTR scores for each protein-coding gene (https://astrazeneca-cgr-publications.github.io/OncMTR-Viewer/).

## Introduction

The availability of large-scale human genetic variation reference datasets has revolutionized our ability to identify disease-causing mutations (Karczewski et al., 2020; Wang et al., 2021). Through the effective process of natural selection, variants with severe clinical outcomes are generally depleted in these datasets. We and others have leveraged this paradigm to develop intolerance metrics that quantify the extent to which natural selection constrains germline variation in genes and genic-sub regions (Dhindsa et al., 2020; Petrovski et al., 2013; Samocha et al., 2014; Traynelis et al., 2017). These methods have proven invaluable in prioritizing which of the roughly 20,000 protein-coding variants observed in any given individual are most likely to contribute to disease. Interpreting variants in the context of cancer suffers from similar challenges as interpreting germline variation: cancer cells often carry thousands of somatic mutations, but only some of these drive the oncogenic process. Despite their success in prioritizing germline variants, population genetics-based approaches have yet to be applied in the context of distinguishing between somatic cancer driving mutations and neutral “passenger” mutations.

Many developmental disorder-causing germline mutations occur in essential genic subregions, leading to dysfunction of crucial cellular biology pathways. We postulated that if these same mutations arise mitotically later in life, they will not cause the same developmental disease due to more limited expression of the mutation but could have equally as profound impacts on cellular biology. Consistent with this, there are several examples whereby identical point mutations that cause severe developmental syndromes when mutated in the germline cause cancer when mutated somatically (Hoischen et al., 2014; Petrovski et al., 2016), including identical mutations in *PTEN, ASXL1* (Hoischen et al., 2011), *EZH2* (Gibson et al., 2012), and others (Kaplanis et al., 2020). Many of these genes are involved in cell proliferation, chromatin remodeling, genome maintenance, and signal transduction pathways. This convergence highlights a subset of genes in the human genome that are crucial to cell biology, whereby disruptive mutations can cause different clinical outcomes depending on their timing, localization, and cellular context.

Here, we hypothesized that regions of genes that are under strong negative selection for germline variation but are exceptionally mitotically mutable would be enriched for variants that increase cancer risk. Identifying germline-constrained but mitotically mutable genic subregions could help prioritize cancer-driving mutations. Here, we focus on missense variants as they are the most observed protein-coding variant class, are becoming increasingly clinically actionable (Hyman et al., 2017), but importantly are also more difficult to interpret than protein-truncating annotated variants. We previously introduced the missense tolerance ratio (MTR), a sliding window-based approach that detects genetic sub-regions depleted of missense variation (Traynelis et al., 2017). In this study, we extended this method to produce a score (OncMTR) to identify germline intolerant but mitotically mutable genic sub-regions by using exome data from 125,748 individuals in GnomAD (Karczewski et al., 2020). We demonstrate that OncMTR effectively predicts driver mutations of hematologic malignancies. We also use 394,694 UK Biobank exomes to illustrate the utility of OncMTR in prioritizing variants in genetic discovery for cancer phenotypes. This work introduces a population genetics approach to identify genic subregions enriched for cancer-related somatic missense mutations.

## Results

### Putative somatic variants in gnomAD

Population-level catalogues of human genetic variation allow for the investigation of selective constraint and mutational patterns in the exome. We used the gnomAD database of 125,748 human exomes to survey both germline and somatic variants (Karczewski et al., 2020). Although the gnomAD variant calling pipeline was tuned to detect germline variation, we reasoned that we may also be able to identify somatic variants that reach a sufficiently high variant allele frequency to be detected through their germline variant caller. Inherited heterozygous germline variants are expected to have an allelic ratio close to 50%. We observed that the distribution of median allelic balance (AB_median) values for gnomAD variants followed a bimodal distribution, with one distribution centered around 50% and another, smaller distribution centered around 20% (**Fig. 1A**).

**Figure 1.**
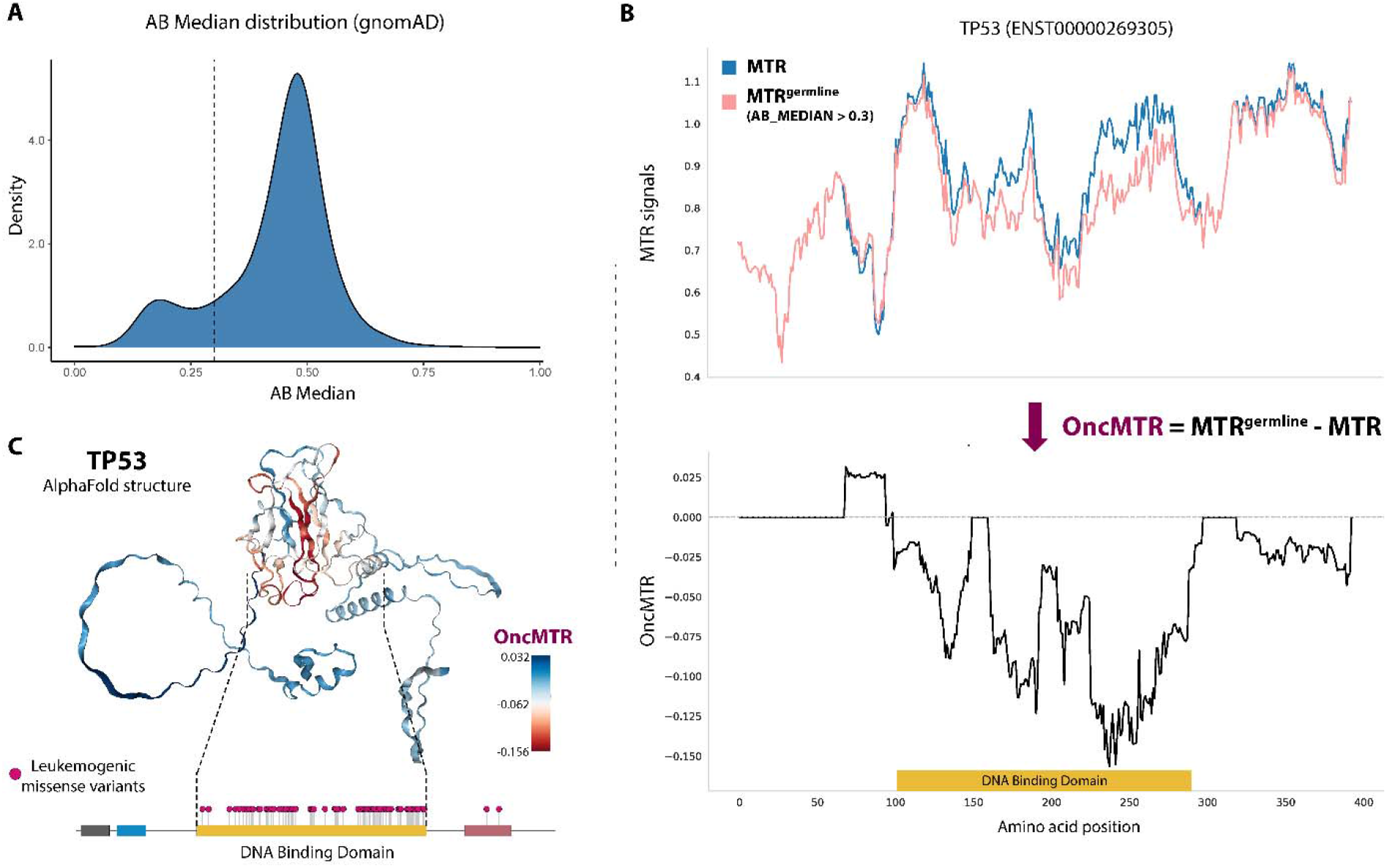
Defining the OncMTR score. **(A)** Bi-modal distribution of median allelic balance values for heterozygous variants in the gnomAD database. We defined putative somatic variants as those with an AB median ≤ 0.3 (dashed line). **(B)** The top figure demonstrates the missense tolerance ratio (MTR) distribution of *TP53* when considering all missense variants (blue) and when restricted to only germline variants (i.e., AB Median > 0.3, depicted in pink). We defined OncMTR as the difference between these two distributions (bottom panel). **(C)** OncMTR scores overlaid on the AlphaFold structure for TP53. The most intolerant region maps to the DNA-binding domain of the protein, which is strongly enriched for mutations known to drive hematologic malignancies.

### Defining OncMTR

We previously introduced a sliding window-based metric, the missense tolerance ratio (MTR), that measures purifying selection on missense variation in genic sub-regions (Traynelis et al., 2017). This score demonstrably detects crucial functional domains of proteins that can cause Mendelian disease when mutated in the germline. Motivated by the overlap between mutations associated with Mendelian disease and cancer, we set out to create a cancer-relevant version of MTR (methods) that captures regions that are depleted of germline variation but also enriched for somatic variation. In this study, we defined another variation of the MTR score, namely MTR^germline^. In its construction, MTR^germline^ is restricted to only those variants achieving an AB_median > 0.3. Taking the well-known cancer gene *TP53* as an example, we can observe those genic subregions where the two MTR formulations diverged (**Fig. 1B**). We then define OncMTR as the difference between these two MTR formulations for each codon and using a 31-codon sliding window (**Fig. 1B**). Negative scores correspond to regions with the greatest divergence between germline intolerance and somatic variant enrichment. Overlaying OncMTR scores on the AlphaFold-predicted structure of TP53 (Jumper et al., 2021) illustrated that the strongest negative scores correspond to the DNA-binding domain, which is the domain enriched for mutations known to drive hematologic malignancies (**Fig. 1C**).

### Using OncMTR to prioritize driver mutations in hematologic malignancies

Motivated by the positive proof-of-concept demonstrated for *TP53*, we next tested whether the MTR and MTR^germline^ distributions differed across other oncogenes included in the Catalogue of Somatic Mutations in Cancer (COSMIC) Cancer Gene Census (CGC). The CGC is divided into two tiers, with Tier 1 containing *bona fide* cancer genes (n=556) and Tier 2 containing genes that have strong indications of playing a role in cancer but with less expansive evidence than Tier 1 (n=137). The difference between MTR and MTR^germline^ distributions per gene, calculated via cross entropy, was significantly higher for Tier 1 genes than a random selection of 556 non-CGC genes (p = 5.7×10^-31^), the remainder of the exome (p = 2.8×10^-67^), and Tier 2 genes (p = 1.1×10^-7^) (**Fig. 2A**). The cross entropy was also significantly larger for Tier 2 genes than the remaining genes in the exome (p = 2.6×10^-4^) (**Fig. 2A**). Together, these results support the hypothesis that mitotically mutable genic sub-regions that are intolerant to germline variation are broadly relevant to cancer.

**Figure 2.**
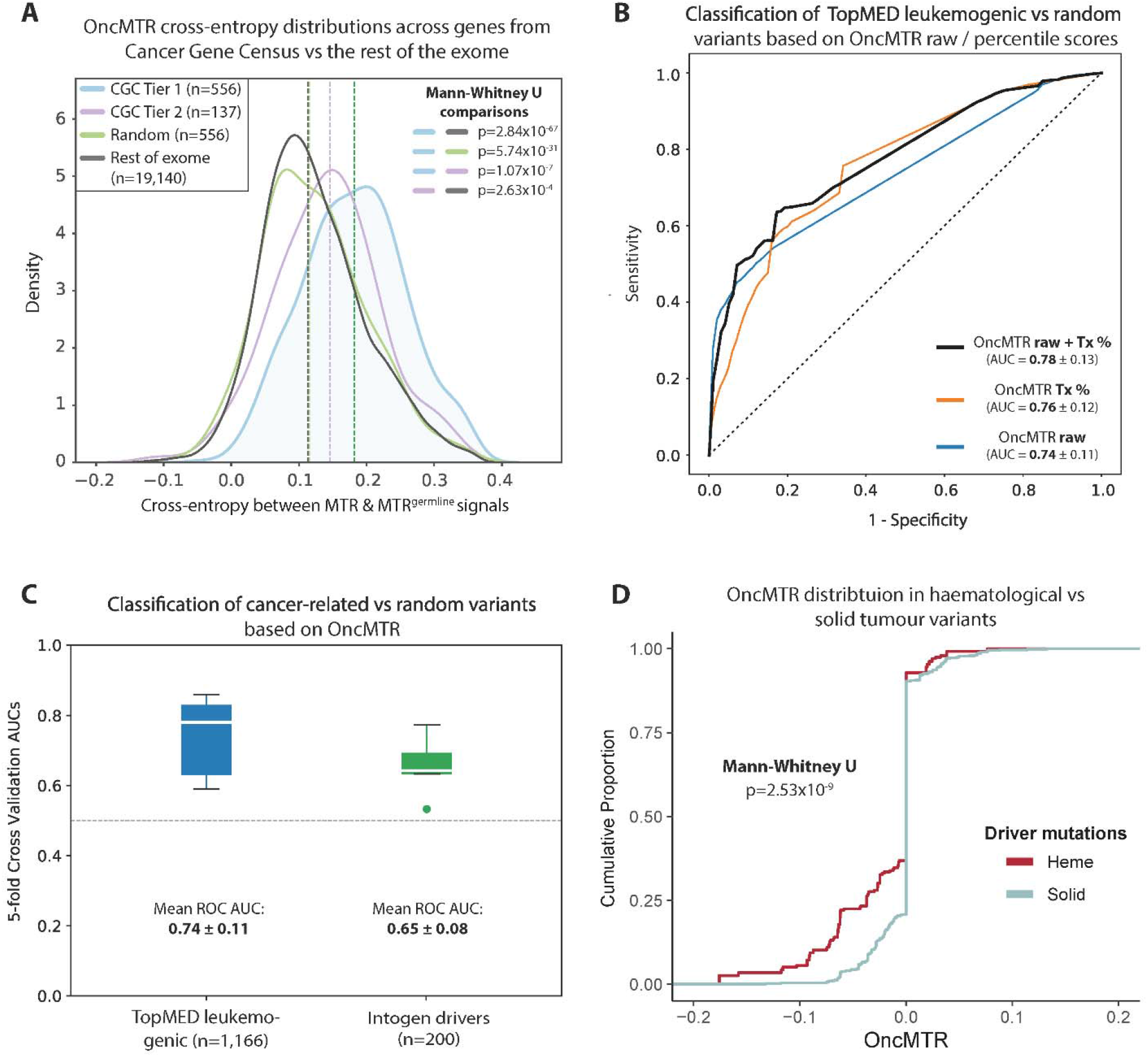
OncMTR regions are enriched for somatic variants associated with hematologic malignancies. (A) Cross entropy between the distribution MTR and MTR^AB^ distributions for Catalogue of Somatic Mutations in Cancer (COSMIC) Cancer Gene Census (CGC) genes, a random selection of genes, and the rest of the exome. **(B)** Receiver operator curve (ROC) depicting the ability of random forest models based on either the raw OncMTR score, the OncMTR transcript-level percentile scores (“Tx%”), and a joint model in discriminating between 1,166 leukemogenic variants and a random size-matched set of variants. AUC = area under the curve. **(C)** Mean ROC AUCs (with fivefold cross-validation) of random forest models based on raw OncMTR in predicting variants involved in leukemia (same variant set as figure B) and hematologic driver mutations annotated in IntoGen (Tamborero et al., 2018). The putatively neutral variant sets comprise of random, size-matched selection of variants. (**D**) The OncMTR distributions of driver mutations for hematologic malignancies versus solid tumors is derived from the Cancer Genome Interpreter.

Distinguishing between cancer-causing driver mutations and neutral passenger mutations remains a central challenge in cancer genomics. We thus tested whether OncMTR could help prioritize somatic mutations that cause hematologic malignancies. We found that the OncMTR scores of a previously defined list of 1,166 leukemogenic driver mutations (Bick et al., 2020) (**Supplementary Table 1**) were significantly lower than a size-matched set of random variants (Mann Whitney U p=2.97×10^-86^; **Supplementary Fig. 1A**). A random forest model using OncMTR achieved an area under the receiving operator curve (AUC) of 0.74 in discriminating between these leukemogenic variants and the random set (**Fig. 2B**). We also calculated transcript-level percentiles for the MTR scores, in which lower percentiles corresponded to lower OncMTR scores. The AUC or the OncMTR transcript percentiles was 0.76, and a combined model that incorporated both the raw OncMTR scores and transcript percentiles achieved an even higher AUC of 0.78 (**Fig. 2B**).

To further assess the capacity of OncMTR to prioritize driver mutations, we trained random forest models with raw OncMTR scores using 5-fold cross-validation. The mean AUC for predicting leukemogenic variants was 0.74 (**Fig. 2C**). We next compared the performance of OncMTR in distinguishing between a set of random variants and 200 established driver mutations implicated in acute lymphocytic leukemia (ALL), acute myeloid leukemia (AML), chronic lymphocytic leukemia (CLL), diffuse large B-cell lymphoma (DLBCL), or multiple myeloma (MM), achieving an AUC of 0.65 (**Fig. 2C**) and having significantly disparate OncMTR distributions from each other (Mann Whitney U p=4.89×10^-5^; **Supplementary Fig. 1B** and **Supplementary Table 2**). Logistic regression-based classifiers achieved similar, albeit marginally lower, AUCs than the random forest models (with AUCs of 0.73 and 0.62 for the two variant sets, respectively), likely due to a small degree of non-linear distribution of OncMTR scores (**Supplementary Fig. 2**). Altogether, these results demonstrate the utility of ou population genetics-based approach in identifying genic sub-regions relevant to hematologic malignancies.

Because the somatic mutations used to calculate OncMTR arose in the blood, we expected that OncMTR would more reliably prioritize driver mutations in hematologic malignancies than in solid tumors. As expected, the OncMTR scores of driver mutations implicated in heme malignancies were significantly lower (Mann Whitney U p=2.53×10^-9^; **Fig. 2D**). To determine whether OncMTR performs better for certain subtypes of heme malignancies, we compared OncMTR distributions of putative driver and passenger mutations identified in a recent comprehensive *in silico* saturation mutagenesis experiment (Muiños et al., 2021). This dataset includes simulated variants across 3 genes for CLL, 9 genes for AML, 2 genes for non-Hodgkin lymphoma, 5 genes for lymphoma, 6 genes for multiple myeloma, and 2 genes for ALL (**Supplementary Table 10**). The OncMTR scores of predicted driver mutations were significantly lower than those of passenger mutations for each cancer subtype, though we observed the strongest separation in CLL (Wilcoxon p<2×10^-308^) and AML (Wilcoxon p=1.4×10^- 155^) (**Supplementary Fig. 3**).

We next assessed whether OncMTR can successfully distinguish between ClinVar pathogenic and benign somatic variants. Logistic regression classification between pathogenic and benign or random variants across all protein-coding genes reached an AUC of 0.60 and 0.58, respectively (**Supplementary Fig. 4**; P=815 unique pathogenic vs B=58 unique benign variants; a set R [random] of equal size to P was sampled to compile the random variants -see also Methods). We next restricted the set of pathogenic somatic variants to those occurring in genes associated with hematologic malignancies and compared to benign or random variants. The AUC was 0.62 in distinguishing between pathogenic and benign variants in hematologic malignancy genes (P=64 vs B=20) and 0.67 when comparing to benign variants across the entire exome (P=64 vs B=58). The AUCs for pathogenic hematologic malignancy variants versus random variants were 0.61 for random variants restricted to heme genes (P=64 vs R=64) and 0.64 for random variants pulled from all protein-coding genes (P=815 vs R=815) (**Supplementary Fig. 4**). These results provide support to this blood-based sequencing version of OncMTR being more powerful in identifying pathogenic mutations implicated with heme malignancies.

Finally, to further explore OncMTR’s power to agnostically detect putative oncogenic regions, we scanned all protein-coding genes in ClinVar in search of transcripts that are preferentially enriched for ClinVar pathogenic somatic variants in regions with OncMTR scores at the bottom 20-percentile of the full OncMTR distribution (see Methods). We identified 101 such transcripts from 24 unique genes (Fisher’s exact test p<0.05; **Supplementary Table 11**), with several known cancer driver genes captured, such as *TP53, IDH1, ALK* and *HNRNPA1* (Martínez-Jiménez et al., 2020). Many of the top ranked genes are implicated in hematologic malignancies, including *MYC, MSH2*, and *FBXW7* (**Supplementary Fig. 5**) (Bhatia et al., 1993; King et al., 2013; Whiteside et al., 2002).

### Genes carrying mutations implicated in both human Mendelian disease and cancer

The underlying hypothesis in deriving OncMTR is that certain genic regions are critically important to human biology, and thus germline mutations in these regions cause severe Mendelian phenotypes, whereas identical somatic mutations–occurring later in life and localized to specific tissue(s)–in these regions may have an oncogenic effect. To evaluate this, we plotted OncMTR distributions for three genes implicated in both neurodevelopmental disease and leukemia: *GNB1, NRAS*, and *DNMT3A* **(Fig. 3 A-C** and **Supplementary Table 4**).

**Figure 3.**
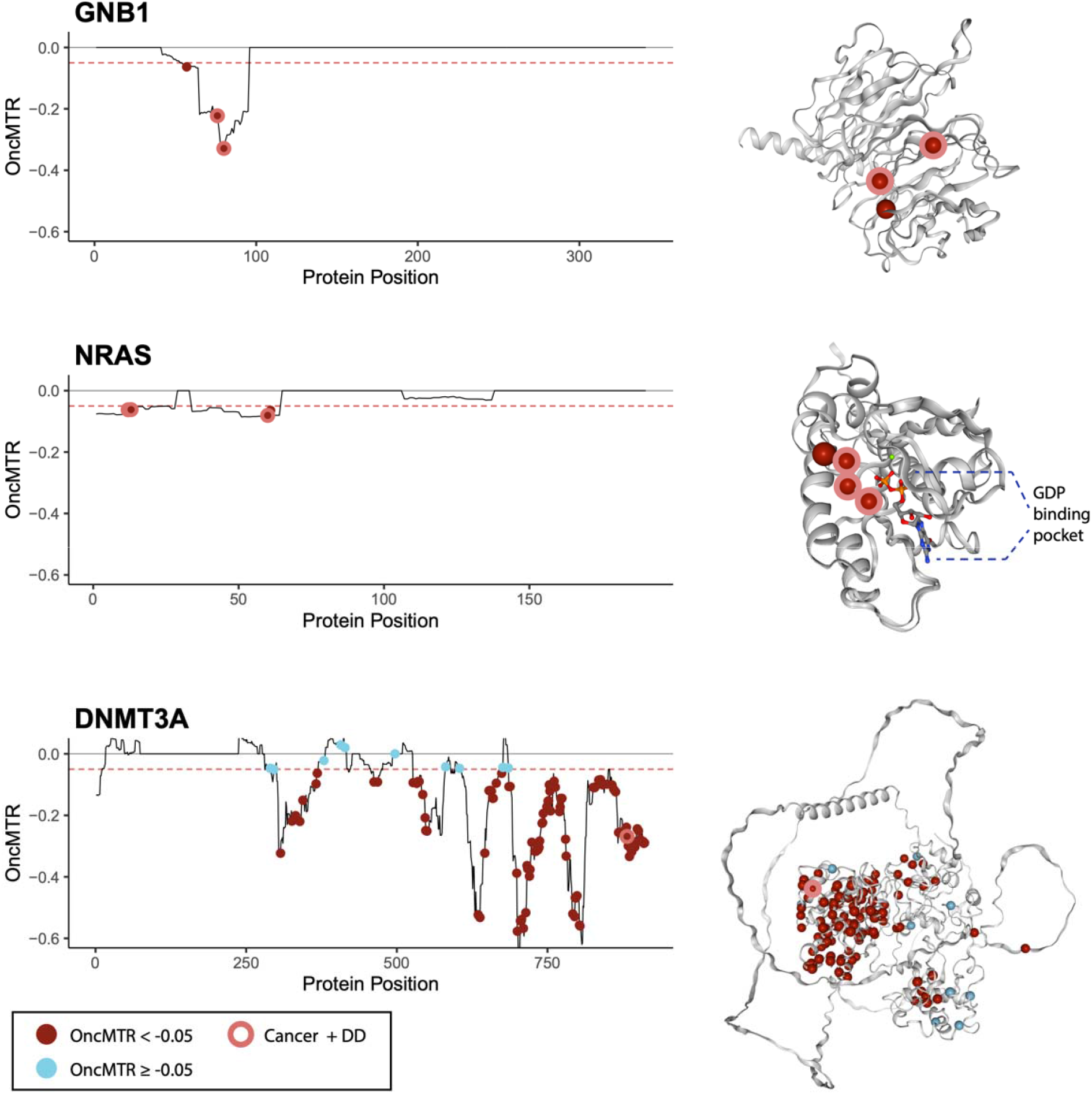
OncMTR distributions for genes implicated in both cancer and Mendelian disease. **(A-C)** OncMTR scores for *GNB1* (A), *NRAS* (B), and *DNMT3A* (C) with corresponding protein structures from PDB (for NRAS, PDB ID: 6zio) or predicted by AlphaFold (Jumper et al., 2021). Points on the OncMTR plots and spheres on the protein structures indicate pathogenic somatic mutations included in TopMED leukemogenic variant set. Red points indicate variants with OncMTR < −0.05. Points with a pink outline indicate somatic leukomogenic variants that are also known to cause developmental delay (DD) when mutated *de novo* in the germline. *De novo* mutations were aggregated from the Online Mendelian Inheritance of Man database.

Germline de novo mutations in *GNB1* cause a severe developmental syndrome characterized by intellectual disability (ID) and other variable features, including hypotonia, seizures, and poor growth (Petrovski et al., 2016). Somatic mutations in this gene have been associated with ALL, CLL, and myelodysplastic syndrome (Yoda et al., 2015). Three of the four somatic driver mutations in this gene overlap with *de novo* mutations implicated in developmental delay (p.Asp76Gly, p.Ile80Thr, and p.Ile80Asn) (**Fig. 3A**) (Petrovski et al., 2016). All four mutated residues reside in a low OncMTR region (OncMTR < −0.05) of the gene, which corresponds to the Gβ-protein surface that interacts with Gα subunits and downstream effectors (**Fig. 3A**).

*NRAS* encodes a RAS protein with intrinsic GTPase activity that has been implicated in multiple hematologic and solid malignancies (Oliveira et al., 2007). There are 28 somatic missense variants in this gene at four distinct amino acid positions associated with juvenile myelomonocytic leukemia and AML, and all residing in low OncMTR regions (**Fig. 3B**) (Bick et al., 2020). Two of these mutations have also been reported as causal germline *de novo* mutations for Noonan syndrome, a developmental delay syndrome that includes congenital heart defects, short stature, and other features (p.Gly13Asp, p.Gly60Glu) (**Fig. 3B**) (Cirstea et al., 2010; Matsuda et al., 2007).

*DNMT3A* encodes a DNA methyltransferase essential for DNA methylation during human embryogenesis and, when mutated somatically, increases risk of acute myeloid leukemia (Kosaki et al., 2017). In a large study on clonal hematopoiesis of indeterminate potential (CHIP), *DNMT3A* was found to harbor the largest proportion of CHIP variants of all CHIP-associated genes (Jaiswal et al., 2017), suggesting it is highly mitotically mutable. In line with this, the OncMTR distribution of this gene is highly enriched for negative values, even compared to *GNB1* and *NRAS* (**Fig. 3C**). The R882 amino acid residue of *DNMT3A* corresponds to a DNA-binding residue that is a major somatic mutation hotspot in CHIP and AML (Kosaki et al., 2017). *De novo* germline mutations at this residue are associated with an overgrowth syndrome called Tatton-Brown-Rahman syndrome characterized by tall stature and impaired intellectual development (Tatton-Brown et al., 2014). Mutations at the R882 residue are thought to interfere with DNA binding, resulting in functional impairment of the protein and aberrant DNA methylation patterns (Zhang et al., 2018). As expected, we identify that the leukemogenic variants in this gene are enriched in low OncMTR regions (**Fig. 3C**). Altogether, these results support the notion that some critically important genic sub-regions are exceptionally mitotically mutable, and mutations in these regions result in different phenotypic outcomes depending on timing and cellular context (Hoischen et al., 2011).

### Enrichment of low OncMTR scores in protein domains

One strength of the sliding window approach implemented in OncMTR is that its estimates are independent of biological boundaries, such as annotated protein domains, which are not always well-annotated. However, it is known that cancer-causing missense mutations tend to cluster in certain functional domains. We thus tested whether Pfam domains and domain superfamilies were enriched for low OncMTR regions (defined as OncMTR < −0.05). Across human protein-coding genes, low OncMTR regions were significantly enriched for several protein domains previously implicated in cancer, such as homeodomains (Fisher’s exact adjusted p-value=4.9×10^-46^), protein kinase domains (Fisher’s exact adjusted p-value=5.25×10^-110^), RING domains (Fisher’s exact adjusted p-value=3.22×10^-48^), and others (**Figure 4 A,B** and **Supplementary Tables 5**,**6**). Furthermore, we found that proteins that had functional domains enriched for low OncMTR scores are significantly enriched in genes with TOPMed leukemogenic variants and known cancer hotspots (Chang et al., 2018) (**Figure 4C** and **Supplementary Tables 1-3;7-9**). Among these two lists of genes, zinc finger motifs were found to be the most strongly enriched for low OncMTR scores (**Figures 4D-F**; most significant adjusted p-value=2.3×10^-52^ from the union list, based on Fisher’s exact test), in line with their well-established role in cancer development (Cassandri et al., 2017). Remarkably, although the calculation of OncMTR is agnostic to domain annotations, it independently identifies cancer-relevant functional genic sub-regions.

**Figure 4.**
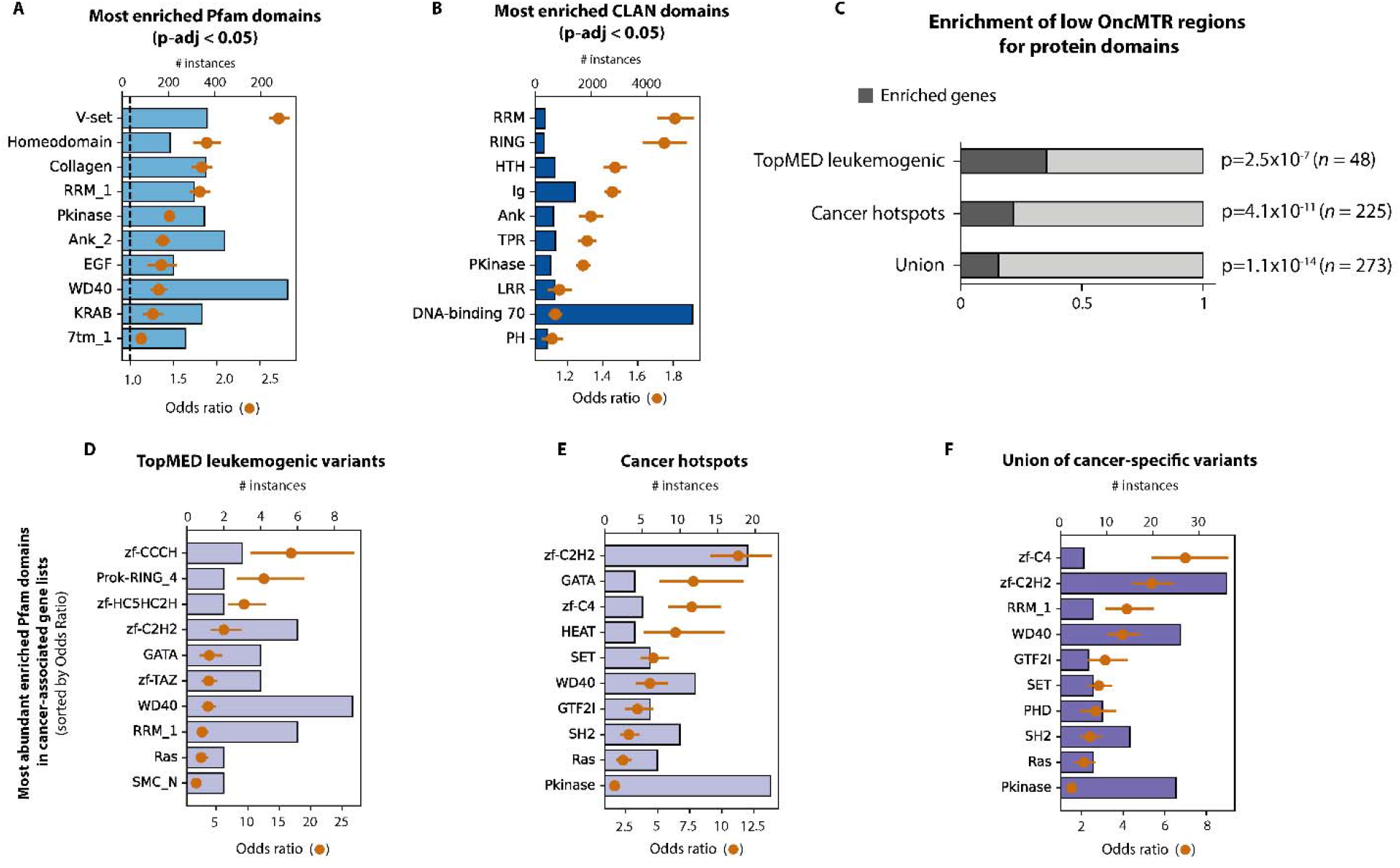
Overlap between OncMTR regions and protein domains. **(A)** Pfam protein domains most strongly enriched with low OncMTR regions (OncMTR < −0.05). **(B)** Pfam domain clans most strongly enriched with low OncMTR regions. The DNA-binding superfamily set was defined in a prior publication (Bahrami et al., 2015). **(C)** Proportions of genes enriched with low OncMTR scores in annotated protein domains in various cancer-related gene sets: genes carrying TopMED leukemogenic variants, annotated cancer hotspots, as well as the union of these three lists. **(D-F)** The most abundant Pfam domains enriched with low OncMTR regions in proteins encoded by the labeled sets of cancer genes. Error bars in each panel represent 95% confidence intervals. P-values were calculated with Fisher’s exact test and adjusted via Bonferroni correction.

### Informing rare-variant collapsing analysis with OncMTR

With increasing adoption of next generation sequencing to generate case-control cohorts, rare variant collapsing analysis has emerged as a powerful approach to detect disease-associated genes for both rare and complex disorders. In this approach, the proportion of cases with a qualifying variant is compared to the proportion of controls with a qualifying variant in the same gene. We have previously shown that incorporating an MTR filter in defining QVs dramatically improves rare variant collapsing analyses (Wang et al., 2021). In that phenome-wide association study (PheWAS) on approximately 300,000 exomes in the UK Biobank, the collapsing analyses detected seven genes associated with hematologic malignancies (Wang et al., 2021). Here, we sought to test whether OncMTR would further improve collapsing analysis signals for hematologic malignancy associations by performing a collapsing analysis on 394,694 European exomes contained in the UK Biobank focused on 1,394 chapter IX (neoplasm) phenotypes. We defined a total of eight collapsing models with and without OncMTR filters (**Supplementary Table 12**). Imposing an OncMTR filter of −0.05 (i.e., only considering missense QVs that fall below this threshold) significantly increased the effect sizes of gene-phenotype associations (p < 0.0001) for each model (**Fig. 5A**). We observed genome-wide significant (p<1×10^-8^) associations between several heme malignancies and *DNMT3A, FBXW7, IDH2, IGLL5, JAK2, SF3B1, SRSF2, TET2*, and *TP53*, in certain cases the effect sizes were 10-fold greater than without adopting the OncMTR filter (**Fig. 5B**). We also found that the association between *TP53* and CLL only reached significance in models including our OncMTR filter; for example, in the ‘raredmg’ model, this association had a p-value of 1.2×10^-7^ (odds ratio [OR] = 8.8; 95% confidence interval [CI]: 4.8-16.0), whereas in the ‘raredmgoncmtr’ model, the same association reached a p-value of 3.4×10^-10^ (OR = 33.2; 95%CI: 16.1-68.7). Thus, applying the OncMTR filter effectively reduces background variation in the setting of gene-level collapsing analysis for haematological malignancy phenotypes and we advise future large-scale haematological malignancy discovery studies to consider adopting OncMTR filter for improved signal detection.

**Figure 5.**
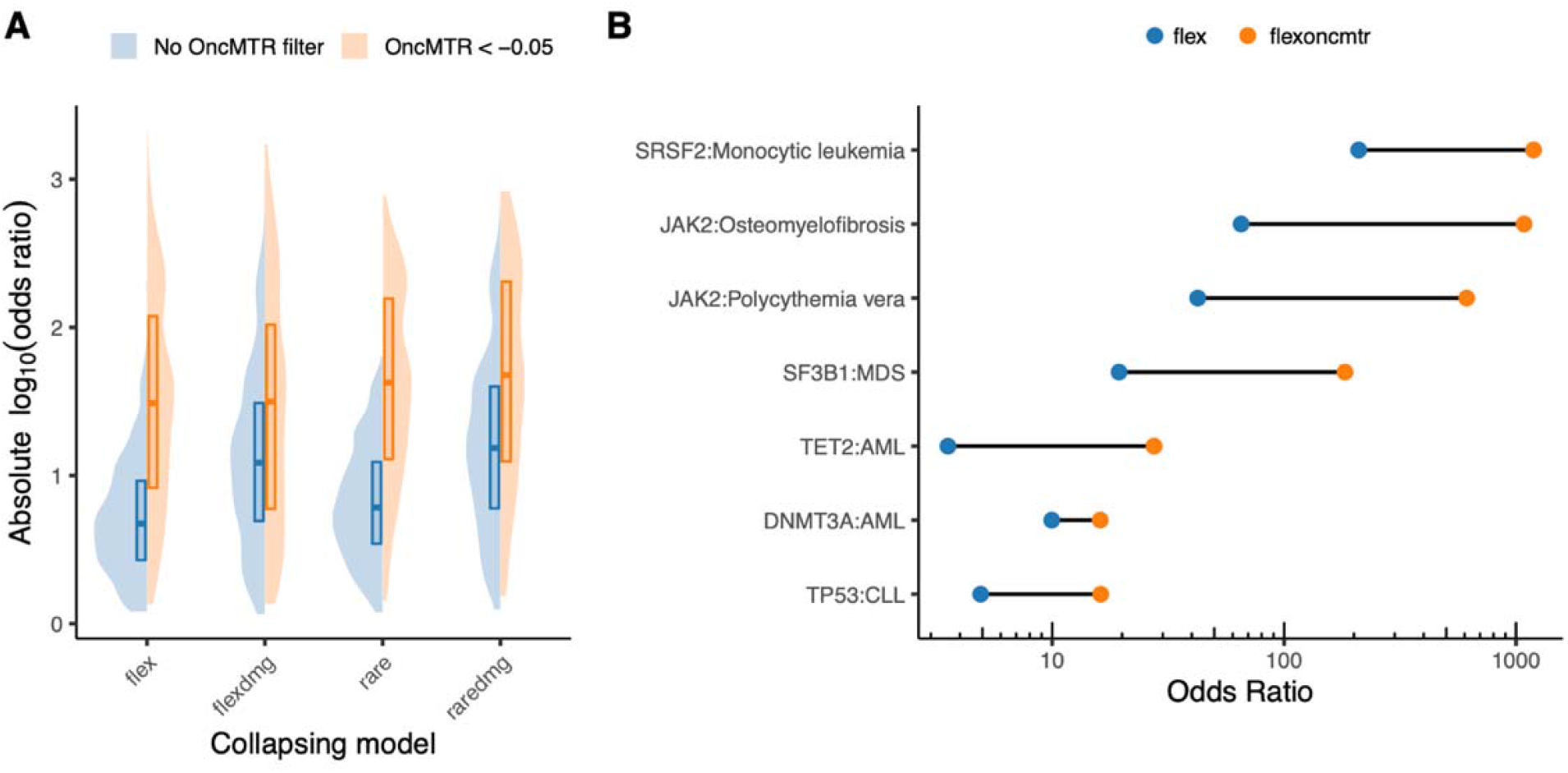
Collapsing analyses using OncMTR. **(A)** Effect sizes of gene-phenotype associations derived from a gene-level collapsing analysis performed on neoplasm phenotypes in 394,694 UK Biobank exomes. Collapsing models are defined in Supplemental Table 12. **(B)** Changes in odds ratios observed for selected gene-phenotype associations. MDS = myelodysplastic syndrome; AML = acute myeloid leukemia; CLL = chronic lymphocytic leukemia.

## Discussion

Determining the clinical relevance of missense variants in oncogenes remains a central challenge in cancer genetics (Chang et al., 2018; Hyman et al., 2017). Motivated by the observation that missense variants in certain genic sub-regions can cause severe Mendelian disease when mutated in the germline and cancer when mutated somatically, we introduced a population genetics-based framework called OncMTR to quantitate the divergence between germline constraint and somatic mutability across the human exome.

First, we demonstrated that oncogenes are enriched for these critically important regions that do not tolerate germline missense variants but harbor somatic mutations. We then illustrated that OncMTR can effectively distinguish between leukemogenic driver mutations and passenger mutations. Although OncMTR is calculated using a sliding window without any input of domain annotations, we found that genic sub-regions that have low OncMTR scores are significantly enriched for protein domains known to be relevant to cancer. Illustrative of our hypothesis was the observation that identical point mutations implicated in both severe Mendelian disease and leukemia in the genes *GNB1, NRAS*, and *DNMT3A* occur in low OncMTR regions. Finally, we found that incorporating OncMTR in a gene-level collapsing analysis on hematologic malignancy phenotypes using 394,694 UKB exomes improved the signal-to-noise ratio for detecting hematologic malignancy associations. We have also developed web server for visualization of OncMTR scores for each human protein-coding gene: https://astrazeneca-cgr-publications.github.io/OncMTR-Viewer/.

Our findings have important implications for the disease biology of both severe Mendelian disorders and cancer. The convergence of genes and genic sub-regions between these two disease areas suggest that similar biological processes play a fundamental role in these two groups of phenotypes. Indeed, cellular proliferation, chromatin remodeling, cell migration, and other cancer-relevant processes have been implicated in neurodevelopmental diseases (De Rubeis et al., 2014; Dhindsa et al., 2021; Feng et al., 2019; Kaplanis et al., 2020). Furthermore, our work supports the notion that mutations in these genes have different phenotypic manifestations based on timing (i.e., zygote versus adulthood), localization (systemic versus hematological), and cellular context.

There exist many other approaches that aim to predict which genic sub-regions are relevant to cancer. These methods tend to look for nonrandom clustering patterns of somatic mutations in either the linear protein sequence or three-dimensional space (Porta-Pardo et al., 2017). To the best of our knowledge, none of these approaches integrate population-level inferences of genic constraint. OncMTR could improve the predictive performance of other, orthogonal driver mutation prediction approaches, as a recent *in silico* saturation mutagenesis experiment demonstrated the strength of incorporating multiple lines of evidence in prioritizing driver mutations (Muiños et al., 2021).

One limitation of OncMTR in its current formulation is that it does not reflect the broader range of solid tumor malignancies since it is based on somatic mutations observed in blood-based sequencing. In future work, the general framework introduced in this study could be applied to sufficiently large tumor-normal sequence datasets as those numbers increase. Furthermore, we used gnomAD because it represents the largest collection of publicly available aggregated allele frequency data. However, gnomAD variants were all called using a germline variant caller. While we demonstrated that we could detect somatic variants in this database, we were likely limited to those that reached a sufficiently high variant allele frequency to be detected. Use of somatic variant callers adopted on these large-scale datasets could further improve the sensitivity of OncMTR.

## Methods

### Reconstructing the Missense Tolerance Ratio with 125K samples from gnomAD

We first reconstruct the Missense Tolerance Ratio (MTR) using a cohort of 125,748 exomes from the gnomAD consortium (v2, GRCh38 liftover). The formula for deriving the window-based MTR scores has been introduced in the original paper (Traynelis et al., 2017):

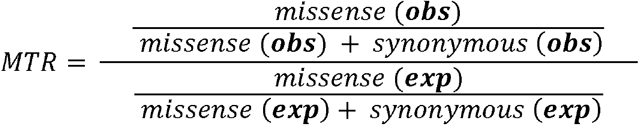

where the numerator represents the observed proportion of missense variants among the total observed protein-coding variation. The numerator is then scaled by the same proportion computed from the collection of all possible protein-coding variants in the corresponding protein-coding window. A window size of 31 codons has been employed for calculating MTR based on the gnomAD cohort, in agreement with the previously published score (Traynelis et al., 2017).

The expected proportion of missense variants in a given protein-coding window was calculated by annotating all possible variants of a protein-coding transcript with SnpEff 4.3t using GRCh38.92 as the reference annotation and assuming all events were equally likely to occur. Annotation with SnpEff focused on single nucleotide variants (SNVs) that were flagged as PASS variants in the original gnomAD release (v2). Variants annotated as ‘missense_variant’ or ‘missense_variant&splice_region_variant’ by SnpEff represent the set of ‘missense’ variants in the MTR formula. Variants annotated as ‘synonymous_variant’, ‘stop_retained_variant’, ‘splice_region_variant&stop_retained_variant’ or ‘splice_region_variant&synonymous_variant’ by SnpEff were considered as the ‘synonymous’ variants in the same formula.

### OncMTR score construction

Using MTR as our basis, we construct the OncMTR score (i.e. Oncology MTR score) to capture protein-coding subregions that are depleted of germline missense variants but observe somatic mutations. We observe that the total distribution of AB_MEDIAN values across all gnomAD variants (**Fig. 1A**) is bimodal, with the main peak centered close to 0.5 and a second one emerging for values approximately around 0.2. The AB_MEDIAN metric represents the allelic ratio between the alleles for each variant, with values close to 0.5 reflecting an equal number of copies being inherited from each parent in heterozygous settings, while truly biological variants that approach zero increasingly reflect variants that more likely arose somatically.

We leverage this observation to construct an alternative version of the original MTR score: excluding any putative somatic variants and employing only germline variants from the gnomAD dataset. We achieve that by selecting only variants with AB_MEDIAN > 0.3, constructing the MTR^germline^ version of the score. OncMTR is then defined as the difference of the original MTR score from the MTR^germline^ version:

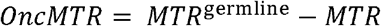

Negative OncMTR values (i.e. MTR^germline^ < MTR) represent regions that are depleted of germline variants and are instead enriched for somatic variation, thus allowing to highlight putative oncogenic subregions in protein coding genes.

### Compilation of variant sets

We used a pre-compiled set of variants known to be drivers of haematologic malignancies in a total of 160 genes (Jaiswal et al., 2014). This list was generated from recurrent haematologic somatic mutations in the literature and COSMIC, excluding genes with a relatively high proportion of loss-of-function germline mutations. A second, smaller pre-compiled list, focused on genes which were recurrent drivers specifically for myeloid malignancies (Bick et al., 2020). A third validation set included a list of annotated driver mutations provided through the IntoGen database (Tamborero et al., 2018). We restricted this set to “Tier 1” (highest confidence) driver mutations observed in hematologic malignancies, which included ALL, AML, CLL, DLBC, and MM.

### Classification of oncogenic variant sets with OncMTR

We have performed classification of different oncogenic variant sets (TOPMed leukemogenic and Intogen drivers) against random variant sets of equal size. We employ two supervised models for the binary classification task: Logistic Regression with ‘max_iter’=1000 and a Random Forest classifier with ‘max_depth’=2, to avoid overfitting on the training set. Each classification was performed as a 5-fold cross-validation task and the mean Area Under Curve (AUC) from all folds is reported to reflect the total average performance of each learning task. The implementations of Logistic Regression and Random Forest were derived from the sklearn Python package (v0.22.1).

We also estimated the optimal OncMTR cut point for each classification by calculating the Youden’s index from each learning task. The average Youden index from all classification tasks performed with Logistic Regression was Y_LR_ = −0.0409 (standard deviation: 0.00126) while for Random Forest it was Y_RF_ = −0.0614 (standard deviation: 0.00057). The mean of the two averages of Youden indexes is −0.05115 or −0.05, after rounding it up to one decimal point for simplicity. We thus consider OncMTR values below −0.05 to have the most distinctive power.

### Identifying OncMTR regions significantly enriched for ClinVar somatic variants

For this analysis, we use all ClinVar somatic variants (ORIGIN=2) from the GRCh38 release (last accessed on 9 June 2019), focusing on those annotated as missense or synonymous. We consider as pathogenic variants those annotated as “Pathogenic” or “Likely_pathogenic” and as benign those annotated as “Benign” or “Likely_benign” (based on ClinVar). Classification between pathogenic and benign (or random) variant sets was performed with a logistic regression classifier with 5-fold cross validation (sklearn, Python package v0.22.1). When restricting the classification to heme-implicated genes, we derived those gene sets based on the Intogen annotation (**Supplementary Table 10**).

In order to identify genes/transcripts across the exome that are preferentially enriched for ClinVar somatic pathogenic variants in regions with low OncMTR scores we employ Fisher’s exact test. Specifically, we scan across each transcript and identify what percentage of the codons in each transcript achieve an OncMTR score at the bottom 20-percentile of the full OncMTR distribution (across the entire transcript). Then, we check whether known pathogenic or likely pathogenic ClinVar missense variants preferentially land in these codons (i.e. corresponding to low OncMTR scores) compared to the rest of the transcript. We apply a Fisher’s exact test (FET) to evaluate the enrichment of each set of regions, i.e., those with low OncMTR scores vs the rest of the transcript. Eventually, we rank each transcript based on the odds ratio and significance of the FET enrichments (**Supplementary Table 11**).

### Enrichment of low OncMTR scores in protein domains

To describe the functional context of OncMTR, we calculated enrichment of constrained regions in protein domain families. Residues within each canonical transcript (as defined by UniProtKB) were divided into two classes based on their corresponding OncMTR scores: below −0.05 (constrained; as defined by Youden’s index) and greater or equal −0.05 (relaxed). Domain and clan annotations for the human proteome were taken from the Pfam 34.0 database. DNA-binding domains were pulled from a previous compendium (Bahrami et al., 2015). The final set of the canonical human proteome consisted of 18,313 annotated proteins. Enrichments of the constrained regions in protein domains were tested with the Fisher’s exact test followed by Bonferroni correction, and significance level of adjusted *p*-value of 0.05.

### UK Biobank Collapsing analysis

Collapsing analyses were performed using the 394,694 exomes available in the UK Biobank (UKB) cohort (Bycroft et al., 2018). The UKB is a prospective study of approximately 500,000 participants aged 40–69 years at time of recruitment. Participants were recruited in the UK between 2006 and 2010 and are continuously followed. The average age at recruitment for sequenced individuals was 56.5 years and 54% of the sequenced cohort is of female genetic sex. Participant data include health records that are periodically updated by the UKB, self-reported survey information, linkage to death and cancer registries, collection of urine and blood biomarkers, imaging data, accelerometer data and various other phenotypic end points. All study participants provided informed consent and the UK Biobank has approval from the North-West Multi-centre Research Ethics Committee (MREC; 11/NW/0382).

We performed a gene-based collapsing analysis on 1,394 chapter IX (neoplasm) phenotypes adopting our previously described approach (Wang et al., 2021). We implemented a total of eight dominant collapsing models with and without OncMTR filters (**Supplementary Table 12**). Using SnpEff (Cingolani et al., 2012), we defined PTVs as variants annotated as exon_loss_variant, frameshift_variant, start_lost, stop_gained, stop_lost, splice_acceptor_variant, splice_donor_variant, gene_fusion, bidirectional_gene_fusion, rare_amino_acid_variant, and transcript_ablation. We defined missense as: missense_variant_splice_region_variant, and missense_variant. Non-Synonymous variants included: exon_loss_variant, frameshift_variant, start_lost, stop_gained, stop_lost, splice_acceptor_variant, splice_donor_variant, gene_fusion, bidirectional_gene_fusion, rare_amino_acid_variant, transcript_ablation, conservative_inframe_deletion, conservative_inframe_insertion, disruptive_inframe_insertion, disruptive_inframe_deletion, missense_variant_splice_region_variant, missense_variant, and protein_altering_variant. We derived allele frequencies from gnomAD (Karczewski et al., 2020). The raredmg, raredmg_OncMTR, flexdmg, and flexdmg_oncMTR models incorporated a REVEL cutoff of REVEL >= 0.5 to restrict to putatively damaging missense variants (Ioannidis et al., 2016).

To compute p-values, the carriers of at least one qualifying variant (QV) in a gene were compared to the non-carriers. The difference in the proportion of cases and controls carrying QVs in a gene was tested using a Fisher’s exact two-sided test. we applied the following quality control filters: minimum coverage 10X; annotation in CCDS transcripts (release 22; approximately 34 Mb); at most 80% alternate reads in homozygous genotypes; percent of alternate reads in heterozygous variants ≤ 0.25 and ≥ 0.8; binomial test of alternate allele proportion departure from 50% in heterozygous state P < 1 × 10^-6^; GQ ≤ 20; FS ≥ 200 (indels) ≥ 60 (SNVs); MQ ≤ 40; QUAL ≤ 30; read position rank sum score ≤ −2; MQRS ≤ −8; DRAGEN variant status = PASS; the variant site achieved ten-fold coverage in ≤ 25% of gnomAD exomes, and if the variant was observed in gnomAD exomes, the variant achieved exome z-score ≤ −2.0 and exome MQ ≤ 30. We excluded 46 genes that we previously found associated with batch effects (Wang et al., 2021).

For all models, we applied the following quality control filters: minimum coverage 10X; annotation in CCDS transcripts (release 22; approximately 34 Mb); at most 80% alternate reads in homozygous genotypes; percent of alternate reads in heterozygous variants ≤ 0.25 and ≥ 0.8; binomial test of alternate allele proportion departure from 50% in heterozygous state P < 1 × 10-6; GQ ≤ 20; FS ≥ 200 (indels) ≥ 60 (SNVs); MQ ≤ 40; QUAL ≤ 30; read position rank sum score ≤ −2; MQRS ≤ −8; DRAGEN variant status = PASS; the variant site achieved ten-fold coverage in ≤ 25% of gnomAD exomes, and if the variant was observed in gnomAD exomes, the variant achieved exome z-score ≤ −2.0 and exome MQ ≤ 30. We excluded 46 genes that we previously found associated with batch effects10.

We defined the study-wide significance threshold as p<1×10^-8^. We have previously shown using an n-of-1 permutation approach and the empirical null synonymous model that this threshold corresponds to a false positive rate of 9 and 2, respectively, out of ∼346.5 million tests for binary traits in the setting of collapsing analysis PheWAS (Wang et al., 2021).

## Acknowledgements

We thank Oliver Backhouse for useful discussions and his feedback on this work. We thank Lawrence Middleton for his assistance in developing the OncMTR web portal.

## Author Contributions

D.V., R.S.D., and S.P. designed the study. D.V., R.S.D., J.M., D.M., J.A., Q.W., B.S., A.R.H., and S.P. performed analyses and statistical interpretation. Q.W. performed bioinformatic processing. D.V., R.S.D., and S.P. wrote the manuscript. D.V. and D.M. developed the web portal. D.V., R.S.D., J.M., D.M., Z.Z., J.A., Q.W., B.S., A.R.H, and S.P. reviewed the manuscript.

## Competing Interests

D.V., R.S.D., J.M., D.M., Z.Z., J.A., Q.W., B.S., A.R.H, and S.P. are current employees and/or stockholders of AstraZeneca.

## Notes

https://astrazeneca-cgr-publications.github.io/OncMTR-Viewer/

