## Supplementary Figures for "Cancer-driving mutations are enriched in genic regions intolerant to germline variation"

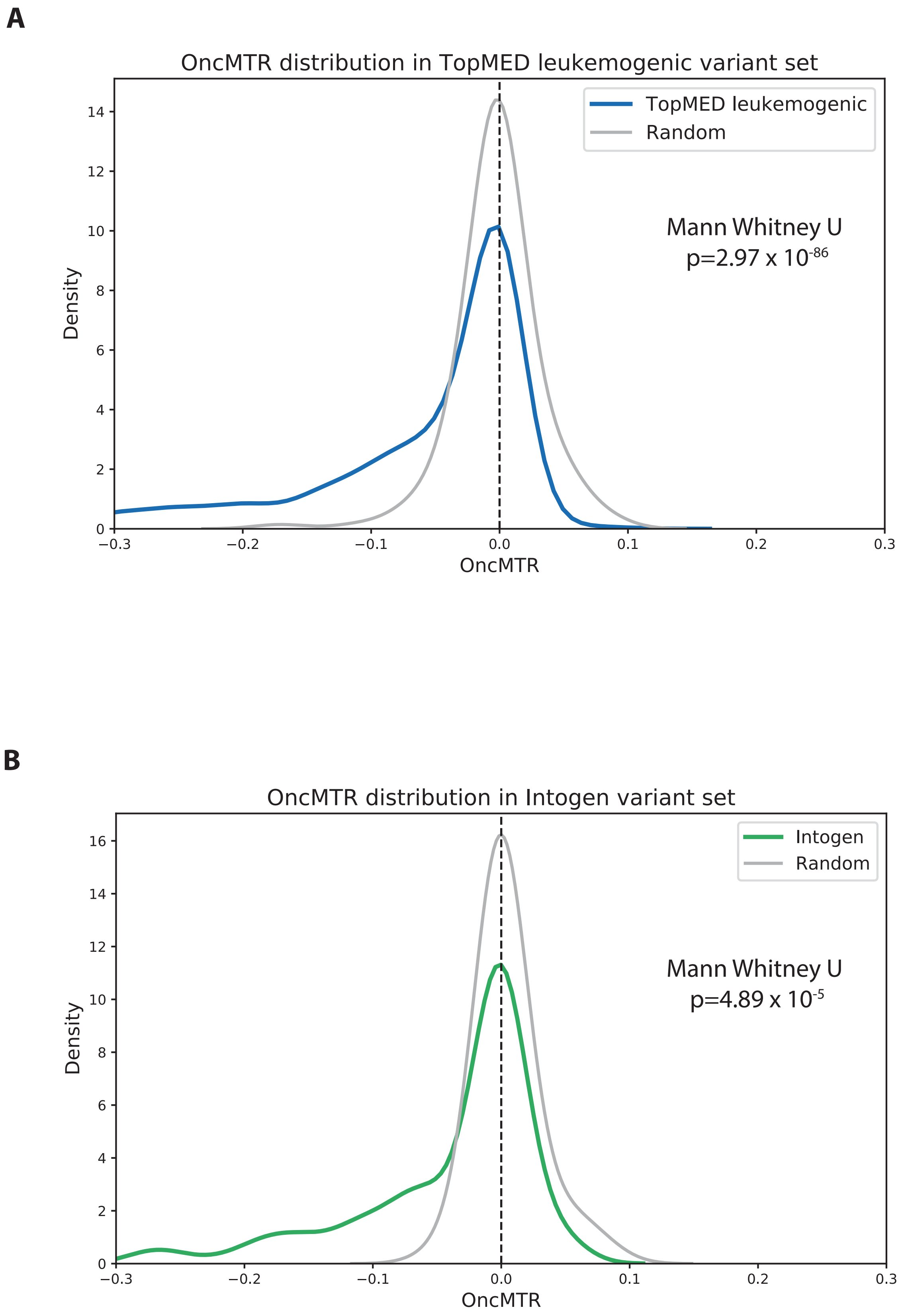


**Supplementary Fig. 1.** OncMTR distributions for cancer-related sets of variants vs size-matched sets of random variants. **A.** TopMED leukomegenic variants. **B.** Intogen driver variants.

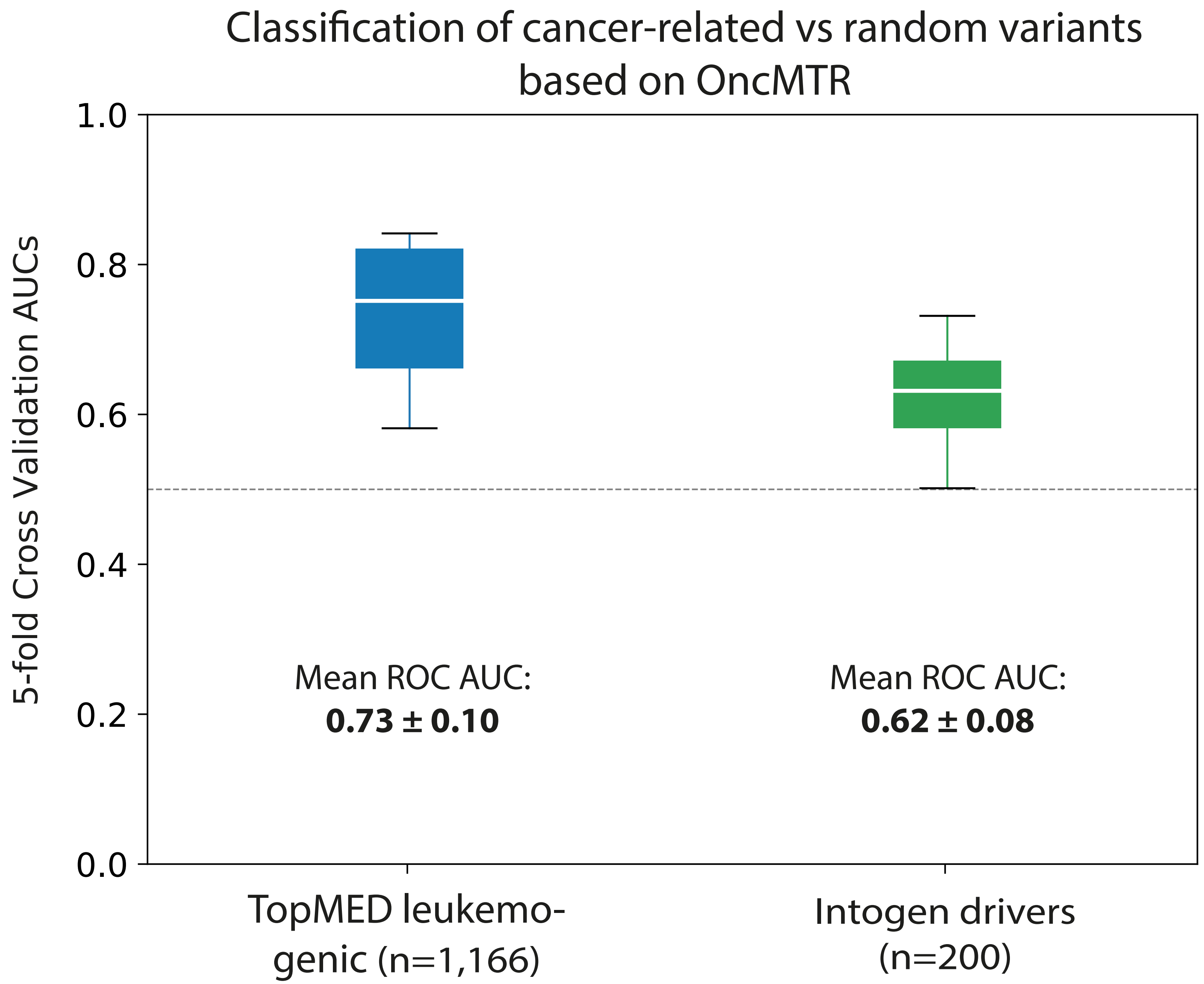


**Supplementary Figure 2.** Mean ROC AUCs (with 5-fold cross-validation) of logistic regression models based on raw OncMTR in predicting variants involved in leukemia (TopMED dataset) and hematologic driver mutations annotated in IntoGen. The putatively neutral variant sets comprise of random, size-matched selection of variants.

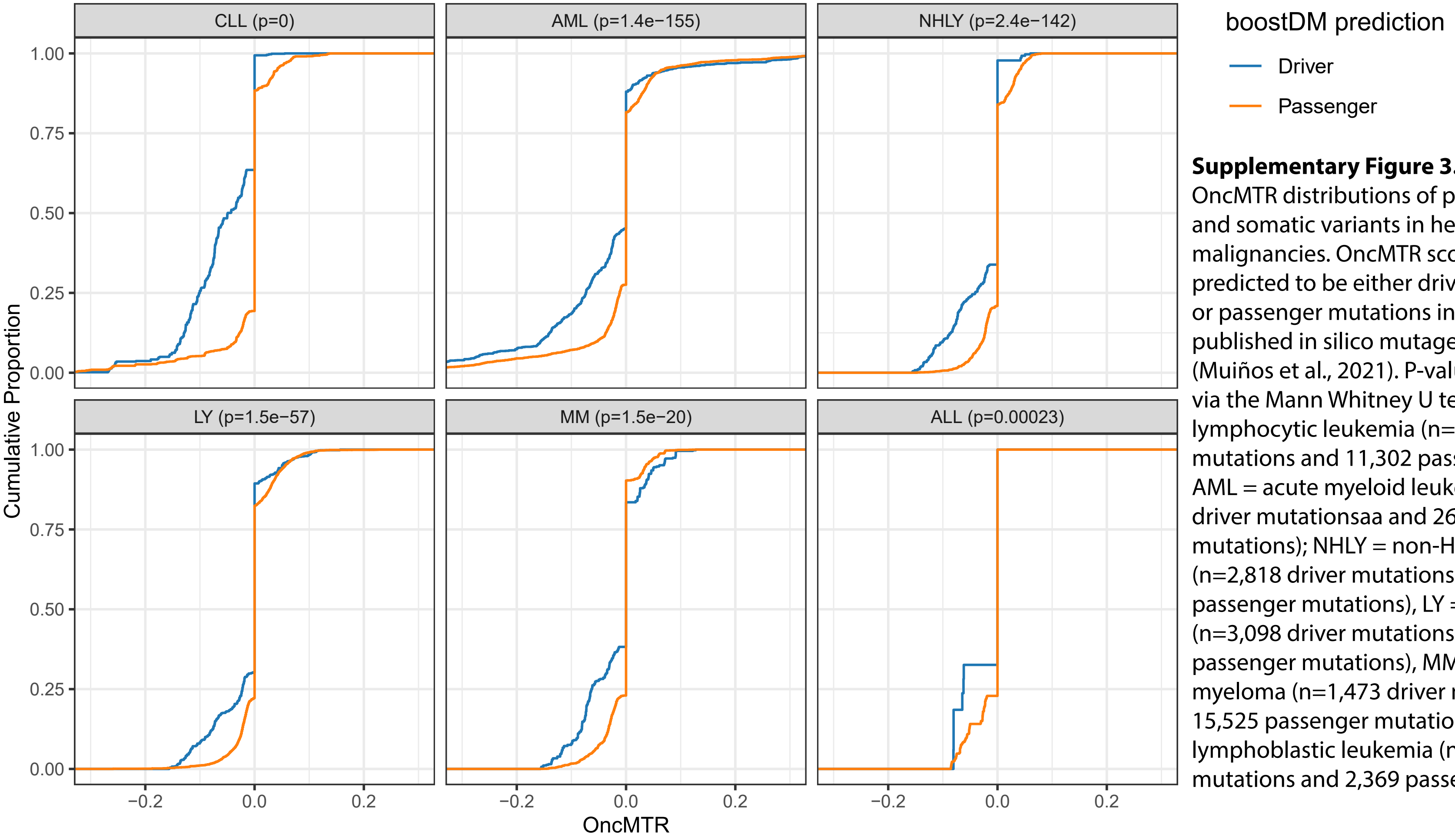


**Supplementary Figure 3.**
OncMTR distributions of putative driver and somatic variants in hematologic malignancies. OncMTR scores of mutations predicted to be either driver mutations or passenger mutations in a previously published in silico mutagenesis experiment (Muiños et al., 2021). P-values were derived via the Mann Whitney U test. CLL = chronic lymphocytic leukemia (n=1,840 driver mutations and 11,302 passenger mutations); AML = acute myeloid leukemia (n=4,795 driver mutationsaa and 26,017 passenger mutations); NHLY = non-Hodgkin lymphoma (n=2,818 driver mutations and 15,841 passenger mutations), LY = lymphoma (n=3,098 driver mutations and 58,459 passenger mutations), MM = multiple myeloma (n=1,473 driver mutations and 15,525 passenger mutations); ALL = acute lymphoblastic leukemia (n=135 driver mutations and 2,369 passenger mutations).

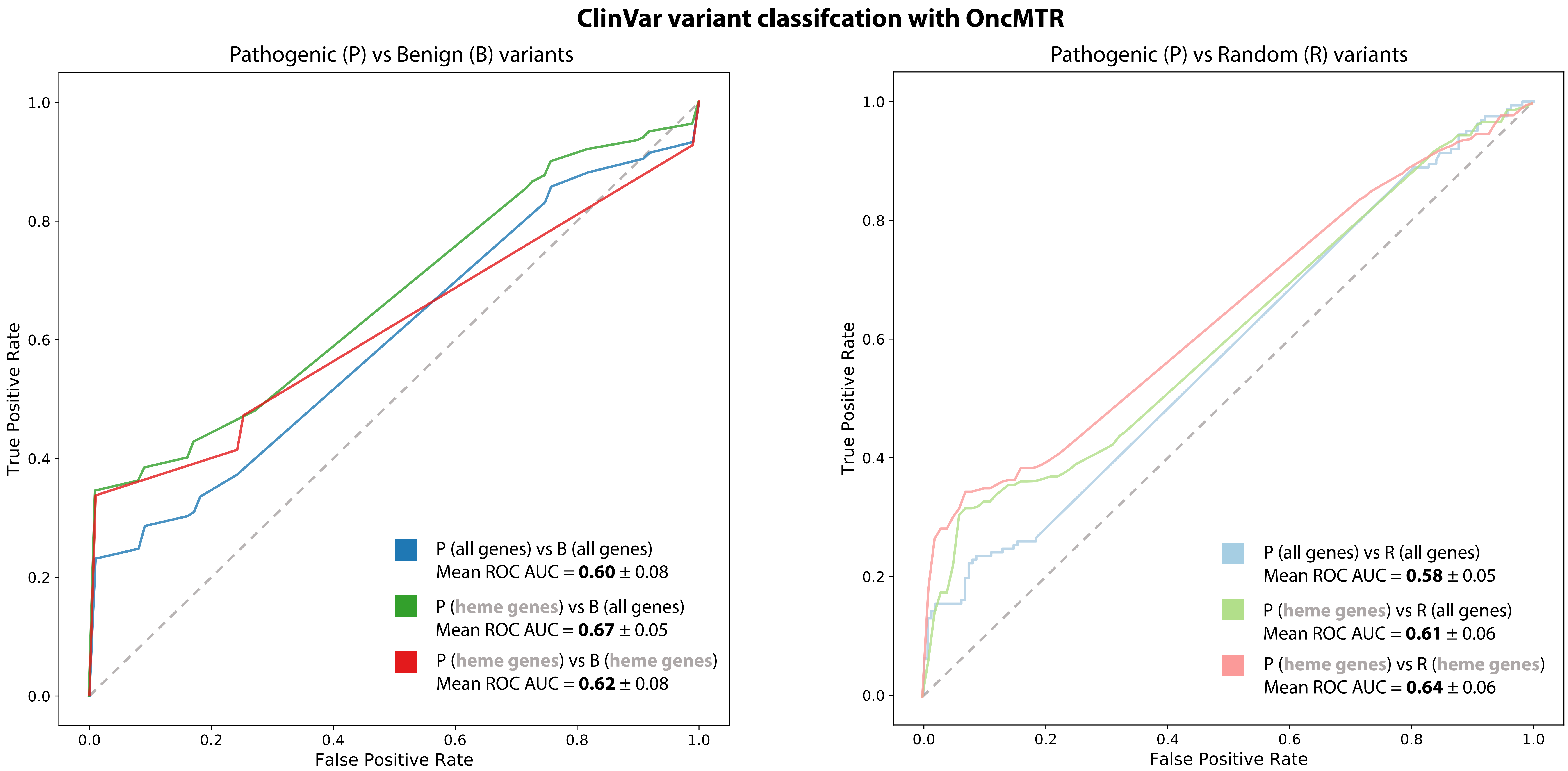


**Supplementary Figure 4.** Mean ROC AUCs (with 5-fold cross-validation) of logistic regression models based on raw OncMTR in predicting ClinVar pathogenic vs benign or random variants. Classification has been performed considering variants across all genes or only on heme genes (noted accordingly in the plots' legend for each case).

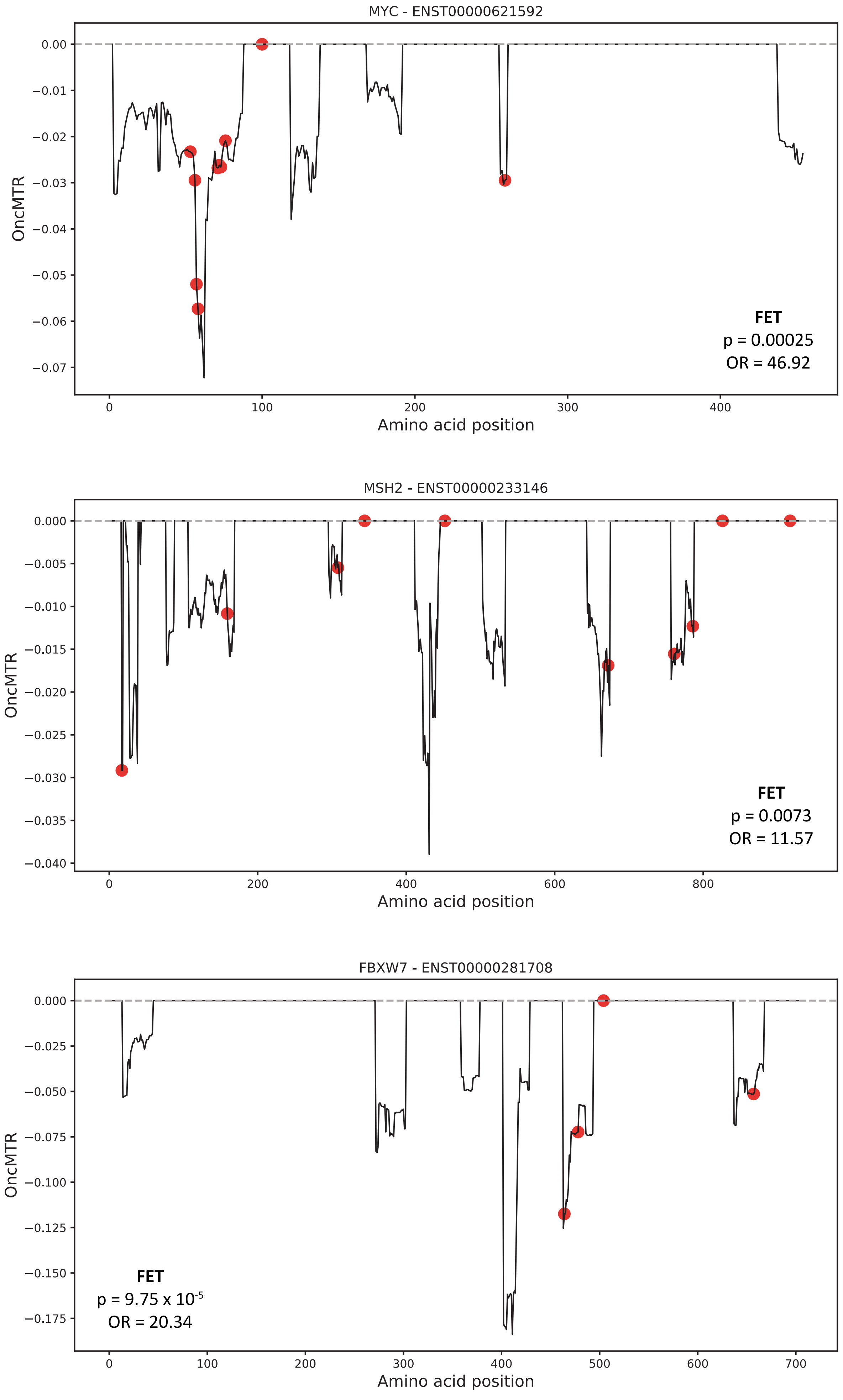


**Supplementary Figure 5.** Example transcripts ranked highly based on the enrichment of ClinVar pathogenic somatic variants falling into regions at the bottom 20-percentile of OncMTR scores vs the rest of the transcript.
